# Generating diffusion MRI scalar maps from T1-weighted images using Reversible GANs

**DOI:** 10.1101/2025.08.27.672677

**Authors:** Tamoghna Chattopadhyay, Gautam Mehendale, Sophia I. Thomopoulos, Himanshu Joshi, Ganesan Venkatasubramanian, John P. John, Jose Luis Ambite, Greg Ver Steeg, Paul M. Thompson

## Abstract

Diffusion tensor imaging (DTI) provides valuable insights into brain tissue microstructure, but acquiring high-quality DTI data is time-intensive and not always feasible. To mitigate data scarcity and enhance accessibility, we investigate the generation of synthetic DTI scalar maps—specifically mean diffusivity (MD)—from structural 3D volumetric T1-weighted brain MRI using a reversible generative adversarial network (RevGAN). Unlike conventional pipelines requiring multiple steps, our approach enables a single-step translation from T1 to diffusion-derived measures. We assess the quality and utility of the synthetic maps in two downstream tasks: sex classification and Alzheimer’s disease classification. Performance comparisons between models trained on real and synthetic DTI maps demonstrate that RevGAN-generated images retain meaningful microstructural features and offer competitive accuracy, underscoring their potential for data augmentation and analysis in neuroimaging workflows. We also examine how well models trained on these data generalize to a new population dataset from India (NIMHANS cohort).

## I. Introduction

Diffusion MRI, particularly diffusion tensor imaging (DTI [1]), is a widely used, non-invasive modality for probing the microstructural organization and connectivity of brain tissue. Despite its utility, acquiring high-resolution diffusion data is resource-intensive, both in terms of scanning time and cost, and not always feasible to collect given time constraints. Moreover, generating reliable diffusion-derived scalar maps requires complex preprocessing—including motion correction, eddy current compensation, and susceptibility distortion correction—followed by model fitting and scalar calculation. Errors at any stage of this pipeline can introduce biases in the final maps. The growing demand for large, high-quality diffusion datasets to train and evaluate machine learning models has intensified interest in data augmentation through synthetic means. Realistically generated diffusion scans have the potential to reduce data scarcity, improve model generalization, and enable robust statistical analyses. Furthermore, synthetic data can alleviate privacy concerns by producing anatomically plausible samples that reflect population-level variability without being linked to any individual subject. In advanced diffusion models such as Mean Apparent Propagator (MAP) MRI [2], even the processing of a single brain slice can require substantial computational time, further underscoring the need for efficient and scalable alternatives like synthetic generation.

While Generative Adversarial Networks (GANs) have demonstrated potential in medical image synthesis, their application to cross-modal MRI translation-particularly between structural T1-weighted images and diffusion-derived scalars-remains underexplored. Recent advances in GAN architectures, such as CycleGAN [3] and conditional GANs (cGANs) [4], have enabled T1-to-T2 translation and FLAIR image synthesis. Even so, these methods face limitations in preserving microstructural features critical for calculating diffusion metrics. Gu et al. [5] used a 2D CycleGAN to generate FA and MD map slices from T1w slices. Pan et al. [6] used a transformer-based encoder along with cGANs to translate 2D T1w to T2w slices. Chattopadhyay et al. [7,8] used denoising diffusion probabilistic models (DDPMs) and latent diffusion models (LDMs) to generate synthetic 2D and 3D DTI-MD maps from T1w MRIs, although they were not one-to-one mapped. Zhang et al. [9] used a diffusion bridge model to generate 3D DTI maps from T1w MRI. We propose a Reversible GAN (RevGAN) [10,11] to address this gap, enabling bidirectional synthesis between T1 images and diffusion scalar maps while maintaining anatomical consistency.

RevGANs extend the traditional GAN architecture by incorporating invertible neural networks. This approach provides several advantages over conventional GANs for our specific task: (1) **Preservation of Information:** The reversible nature of our architecture ensures that structural information present in T1-weighted images is preserved during the translation process, which is crucial for generating accurate diffusion metrics. (2) **Cycle Consistency without Additional Networks:** Unlike CycleGAN, which requires separate generators for forward and backward transformations, RevGANs achieve cycle consistency through a single invertible network, reducing model complexity and training instability.

We analyzed the quality, fidelity, and diversity of data generated from the model, and its effectiveness for supporting downstream classification tasks, offering insights into the practical utility of the model.

## II. Imaging Data and Preprocessing

The primary dataset for our experiments was the widely-used, publicly available Alzheimer’s Disease Neuroimaging Initiative (ADNI) dataset – a multisite study launched in 2004 to improve clinical trials for the prevention and treatment of AD [12]. We used 930 scans for our analysis. The second, held-out dataset comes from an Indian population assessed at NIMHANS in Bangalore, India [13,14,15] – a population not typically well represented in neuroimaging studies. This cohort had 226 participants. The data distribution is shown in **Table 1**; we ensured that the testing and training data subsets did not overlap, and the test dataset had only one scan per subject. 3D T1-weighted (T1w) brain MRI volumes were pre-processed using the following steps [16]: nonparametric intensity normalization (N4 bias field correction), ‘skull stripping’ for brain extraction, nonlinear registration to an in-house template [16] with 6 degrees of freedom and isometric voxel resampling to 2 mm. The pre-processed images were of size 91×109×91. The T1w images were scaled using min-max scaling to take values between 0 and 1. All T1w images were aligned to the ENIGMA template provided by the ENIGMA consortium [17].

**Table 1.** Data distribution for experiments.

| Dataset | | Total N | Sex | | Mean Age ± St. Dev. | Diagnosis | | |
| --- | --- | --- | --- | --- | --- | --- | --- | --- |
|  |  |  | Male | Female |  | CN | MCI | Dementia |
| ADNI | Training Dataset | 805 | 395 | 410 | 73.6 ± 7.8 | 456 | 275 | 74 |
| | Test Dataset | 125 | 66 | 59 | 73.9 ± 8.2 | 41 | 42 | 42 |
| NIMHANS | Test Dataset | 226 | 125 | 101 | 66.9 ± 7.9 | 128 | - | 98 |

**Table 2.**
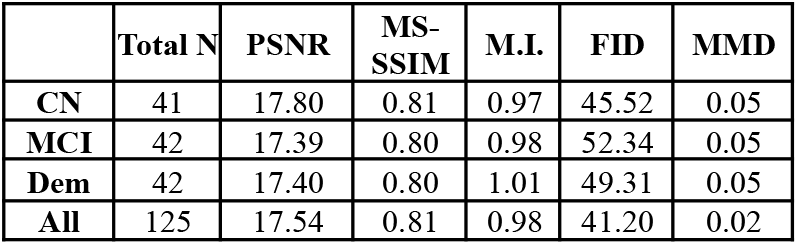
Qualitative Metrics for the Test Dataset. CN indicates healthy control subjects, MCI indicates Mild Cognitive Impairment and Dem indicates Dementia. All indicates all subjects combined. M.I. indicates Mutual Information.

## III. Deep Learning Architectures

The 3D RevGAN architecture (**Fig. 1**) converts T1w scans to synthetic DTI-MD scalar maps using a generator and two discriminators, trained adversarially to enhance realism and structural fidelity of the generated images. The generator transforms 3D T1-weighted inputs of size 91×109×91 into synthetic MD maps of the same dimensions. The network begins with two 3D convolutional layers, each using 3×3×3 kernels with stride 2, instance normalization, LeakyReLU activation, and dropout (0.2). These layers progressively reduce the spatial resolution while expanding feature depth. The encoded features pass through a core component consisting of *R* repeated 3D reversible blocks, each incorporating residual connections and 3×3×3 convolutions with LeakyReLU and dropout. The decoder then reconstructs the output using two transposed convolutional layers. The first upscales the feature maps from 24 to 12 channels, while the final layer reduces the channel count to 1 with a *tanh* activation. The decoder also uses LeakyReLU and a smaller dropout (0.1). This design restores image resolution while preserving learned structural information. The discriminator comprises two identical sub-networks (D1 and D2); both employ 3D CNNs to distinguish real from synthetic image pairs. D1 assesses the realism of (Real T1, Synthetic T1) pairs, while D2 evaluates (Synthetic MD, Real MD) pairs. Each discriminator operates on input tensors of shape 2×91×109×91, passing through three convolutional layers with kernel size 3×3×3 and padding to maintain spatial resolution. The first two layers use LeakyReLU and instance normalization, while the final layer uses a sigmoid activation to produce a probability map for real/fake classification. The network is trained using the Adam optimizer with parameters β_1_ =0.5, β_2_ =0.999, a learning rate of 1×10^−5^, and a batch size of 1 for 600 epochs. Learning rate decay is applied every 200 epochs. Weights are initialized using a normal distribution, *N*(0,0.02).

**Figure 1.**
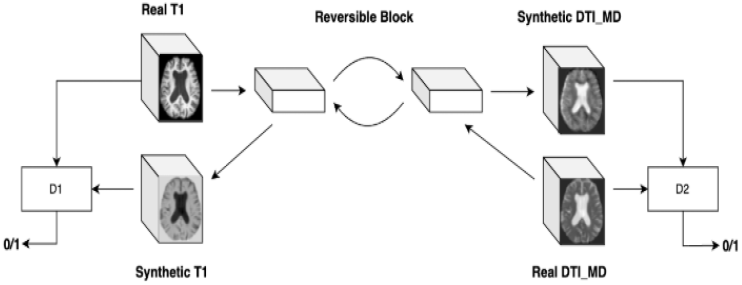
RevGAN architecture.

The 3D CNN model architecture (**Fig. 2**) comprised four convolutional layers with 3×3 kernels, followed by a 1×1 convolution and a final fully connected layer. ReLU activation and Instance Normalization were applied throughout to enhance training stability. To address overfitting, dropout layers with a 0.5 rate were added after the second, third, and fourth convolutional layers, along with a 3D average pooling layer using a 2×2 kernel. The model was trained with the Adam optimizer [18], initialized with a learning rate of 1×10^−4^ and an exponential decay factor of 0.96. Training was performed over 100 epochs with a batch size of 8, optimizing for mean squared error. Early stopping and dropout were used as regularization strategies. Hyperparameters were tuned to identify the optimal configuration. Model evaluation was conducted on two tasks: sex classification (a standard benchmark with known ground truth) and dementia classification, using balanced accuracy, F1 score and AUC-ROC as performance metrics. The NIMHANS dataset served as the independent holdout test set for both tasks. Model performance was compared between original DTI-MD and synthetic DTI-MD maps.

**Figure 2.**
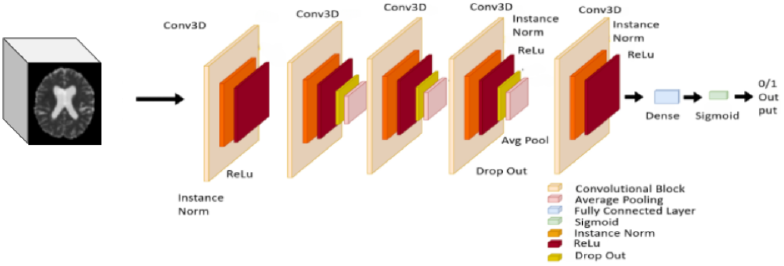
3D CNN Architecture that we trained for downstream tasks.

**Table 3.** Classification results for the synthetic and holdout test sets. B.A. indicates balanced accuracy, F1 indicates F1 Score and A.R. indicates AUC-ROC Scores. The NIMHANS dataset is treated as a zero-shot test dataset.

| Trained On | Tested On | Sex Classification |  |  | Dementia Classification |  |  |
| --- | --- | --- | --- | --- | --- | --- | --- |
|  |  | B.A. | F1 | A.R. | B.A. | F1 | A.R. |
| Original DTI | Original DTI | 0.79 | 0.76 | 0.86 | 0.82 | 0.77 | 0.85 |
| Original DTI | NIMHANS | 0.76 | 0.73 | 0.82 | 0.82 | 0.79 | 0.90 |
| Synthetic DTI | Synthetic DTI | 0.82 | 0.80 | 0.89 | 0.82 | 0.77 | 0.86 |
| Synthetic DTI | Original DTI | 0.78 | 0.78 | 0.82 | 0.77 | 0.70 | 0.79 |
| Synthetic DTI | NIMHANS | 0.76 | 0.78 | 0.83 | 0.79 | 0.77 | 0.88 |

## IV. Results

The 3D RevGAN model architecture generated synthetic DTI-MD scans for all subjects in the test set (**Fig. 3**). To evaluate the quality of synthetic mean diffusivity (MD) maps generated from T1-weighted MRI scans using the RevGAN architecture, we computed a comprehensive set of metrics across clinical subgroups—Cognitively Normal (CN), Mild Cognitive Impairment (MCI), and Dementia (Dem)—from the test dataset. Quantitative evaluations included Peak Signal-to-Noise Ratio (PSNR), Multi-Scale Structural Similarity (MS-SSIM), Mutual Information, Fréchet Inception Distance (FID), and Maximum Mean Discrepancy (MMD) [19].

**Figure 3.**
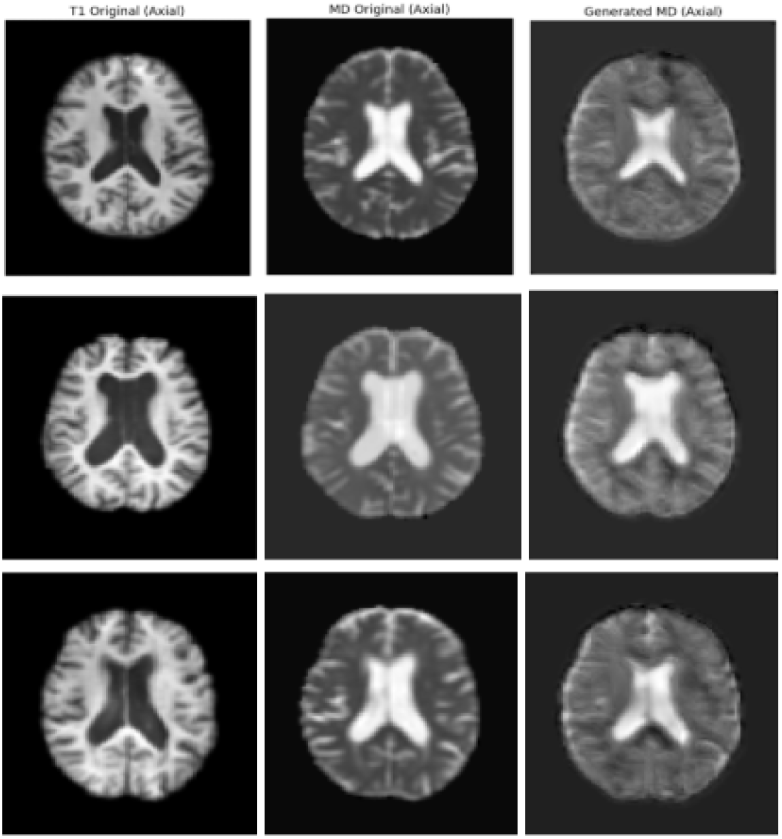
Results from our RevGAN implementation. Each column shows the middle slice from the original T1w scan, the original DTI-MD scan and the synthetic DTI-MD scan respectively. The first row shows a randomly-selected healthy control (CN) subject, the second row shows an MCI subject and the third row shows subjects with Dementia.

The average PSNR across all subjects was 17.54. Subgroup scores slightly varied, potentially reflecting anatomical and structural variability in advanced disease. MS-SSIM scores (∼0.80–0.81) indicate moderately preserved structural similarity between real and synthetic MD images. Notably, mutual information increased from CN to Dem (0.97 to 1.01), suggesting stronger statistical dependence between T1w and MD images in subjects with greater neurodegeneration. FID and MMD, which assess distributional similarity, were lowest for the combined dataset (41.20 and 0.02 respectively), indicating improved generative performance when learning from the full data distribution. Higher FID for individual groups, especially MCI (52.34), may reflect greater inter-subject heterogeneity and transitional pathology that complicates image synthesis. Overall, the results suggest that RevGAN captures global structural features while being sensitive to underlying disease-specific anatomical variability.

To assess the utility of the synthetic DTI maps for downstream predictive modeling, we compared the performance of the 3D CNN architecture for two tasks - Sex Classification (a commonly used benchmark, as ground truth is known) and Dementia Classification. The model trained on synthetic DTI achieved performance closely matching that of the model trained on original DTI, with only marginal differences across metrics. For sex classification, synthetic DTI even outperformed original DTI in balanced accuracy (0.82 vs. 0.79), F1 score (0.80 vs. 0.76), and AUC-ROC (0.89 vs. 0.86), suggesting that the RevGAN-generated images preserved biologically meaningful features relevant for sex prediction. AD classification performance was nearly identical between original and synthetic DTI, further supporting the fidelity of the synthetic data. In zero-shot evaluation on the independent NIMHANS dataset, both tasks showed a minor drop in performance, likely due to domain shift between the ADNI training set and the external test distribution. Notably, synthetic DTI generalized comparably to original DTI in this setting, reinforcing its potential for broader clinical and research applications. The close alignment of classification performance across real and synthetic modalities validates the representational quality of the generated scans and indicates their suitability as substitutes in resource-constrained or privacy-sensitive scenarios.

## V. Conclusions and Future Work

In this study, we demonstrated the efficacy of a RevGAN-based architecture for translating T1-weighted brain MRI scans into synthetic diffusion MRI (DTI-MD) maps using the ADNI dataset. Quantitative image similarity metrics indicated that the synthetic MD maps preserved structural and distributional properties across clinical subgroups, with strong mutual information and favorable FID/MMD scores, particularly when trained across the full dataset. Importantly, downstream evaluation using a 3D CNN on sex and dementia classification tasks revealed that models trained on synthetic DTI performed comparably to those trained on original DTI, and retained generalizability when tested in a zero-shot manner on the independent NIMHANS dataset. These findings validate the clinical relevance of the generated images and highlight the potential of generative approaches in reducing the reliance on hard-to-acquire diffusion scans. Future work will focus on improving fine-grained anatomical fidelity using attention-based or diffusion-based architectures. We will also extend the framework to multi-channel DTI synthesis (e.g., FA, RD, AD, and multishell metrics, e.g. NODDI) for richer downstream utility. Another interesting challenge is scalar-to-tensor translation, or inferring the full diffusion ODF (orientation density function) field from T1w data. Additionally, domain adaptation strategies could further enhance robustness across sites and scanners, supporting real-world deployment in diverse clinical settings.

## Acknowledgments

This work was supported by NIH grant R01 AG060610 funded by the National Institute on Aging (NIA) and the Fogarty International Center (FIC), as well as by NIH NIA grant U01 AG068057 (‘AI4AD’). NIMHANS data collection was supported by the Department of Science and Technology, Govt. of India, grant nos. DST-SR/CSI/73/2011 (G), DST-SR/CSI/70/2011 (G) and DST/CSRI/2018/249 (G).

